# Cognitive resilience in bilinguals and its potentiality in dealing with cognitive load

**DOI:** 10.1101/2020.09.17.301606

**Authors:** Setareh Doroud, Zari Saeedi, Narges Radman

## Abstract

Previous studies have demonstrated different patterns of results regarding cognitive benefits of bilingualism, ranging from bilingual advantage to no effect of bilingualism. This study examined the potential effect of bilingualism on cognitive resilience and performance. We recruited 21 Persian monolinguals and 19 Persian-English bilinguals. Color-Word Stroop task was used concurrently with verbal production tasks in order to produce three levels of task difficulty, i.e., doing the Stroop task while being silent (level 1), alphabet reciting (level 2), and counting odd numbers (level 3). We investigated the pattern of changes in Stroop task performance when faced with different difficulty levels Bilinguals showed less change in their performance in the Stroop task when faced with high cognitive load (high task difficulty level). However, monolinguals showed a significant decrease in their performance when the cognitive load increased. Our data support the “Bilingual Advantage” view. However, this advantage is highlighted in cognitively demanding tasks.

## Introduction

### Cognitive load

As instigated by Sweller, Cognitive Load Theory (CLT) clarifies that handling more complex new information occupies more space in our limited working memory (WM), consequently, performing such tasks would be hindered (Sweller, 1988). In fact, because WM is capacitively limited and keeping the information in WM is temporally short, the individual would be cognitively overloaded in case of processing complex tasks and storing chunks of information(Chen et al., 2012).

Cognitive load is divided into three subcomponents, intrinsic, extraneous, and germane load (Paas et al., 2003; Sweller, 1994). Intrinsic load is defined as the “intrinsic complexity of the core information” (i.e., the level of interactivity or connectedness between different elements of a task to be learned) (Seufert et al., 2007; Sweller, 1994; Windell and Weibe, 2007). The extraneous load is described as the extra load in connection with how the information is presented (Jong, 2010; Windell and Weibe, 2007). Germane load refers to the mental resource devoted to schematic automation of the information stored in long-term memory (Debue and Van de Leemput, 2014). Accordingly, as Sweller asserted (Sweller, 2010), this type of cognitive load is based on the characteristics of the individuals including motivation and cognitive ability, and it does not rely on task characteristics.

### Measuring Cognitive Load

Cognitive load has been specified by an increase of the load on cognitive, physical, and neural aspects of the human body and brain. Subjective measurement of the cognitive load consists of self-report rating scales in which the difficulty of a particular task should be estimated. The physiological effects of the increased cognitive load have been investigated using measures of autonomic nervous system activity (e.g., heart rate, respiratory rate, pupil size) (Brunken et al., 2004; Skulmowski and Rey, 2017). An objective measure of cognitive load is obtained by assessing a participant’s reaction time in response to a task, while attention is also devoted to a secondary task concurrently (Skulmowski and Rey, 2017). An example of such a task is the Stroop task which has been widely used to measure cognitive load (Block, 2004; Gwizdka, 2010; Hazan-Liran and Miller, 2017; Lavie and Viding, 2004).

In addition to behavioral tests, neuroimaging methods have rarely been used to assess cognitive load; namely, Kumar and Kumar (Kumar and Kumar, 2016), testing a group of 6 participants, used power spectrum analysis to assess cognitive load in Human-Computer Interaction. The authors reported an improvement in alpha and beta band powers in response to an increased load of WM (Kumar and Kumar, 2016). One recent work reported that increased cognitive load affects susceptibility to auditory signals. The authors suggest that cognitive load can reduce susceptibility to auditory signals and alerts in an auditory oddball task. This effect was seen by a reduced P300 amplitude when the cognitive load increased (van der Heiden, R. M., Janssen et al., 2020).

### Cognitive resilience

When a high cognitive load is imposed, a factor affecting the performance in both daily life and professional activities is cognitive resilience. Cognitive resilience is the capacity to maintain cognitive ability at risk of loss, depletion, or impairment when facing challenges of high demand (Jha et al., 2017). Resilience has been defined not only as recovery from stress to normal levels of mental health, but also as sustained growth due to the result of healthy responses to stressful situations (Gu and Day, 2013; Reich et al., 2011; Wei et al., 2011). People vary in terms of psychological resilience and cognitive resilience because of variations in cognitive capacity to react to stressful and (Parsons et al., 2016; Stine-Morrow and Chui, 2012).

While cognitive resilience has been measured using self-questionn(Bucur, 2015; Ram et al., 2019), few studies have also applied cognitive tasks with different difficulty level to evaluate how subjects manage the effect of task difficulty at behavioral level (Shields et al., 2017).

### Cognitive resilience in bilingual individuals

One of the factors which mediate cognitive resilience is bilingualism. A great body of research suggests that bilinguals outperform monolinguals in cognitive performances. This assumption is originated and is based on the idea that bilinguals have the ability to inhibit the non-target competing language on a daily basis (Bialystok et al., 2012). Several factors have been introduced to mediate this effect such as the age of learning the second language (Bialystok and Poarch, 2014; Birdsong, 2018; Liu et al., 2017). However, the superiority of bilinguals in cognitive performance has been recently challenged by several authors such as Paap and colleagues (Paap et al., 2016; Paap and Greenberg, 2013) as well as Ratiu and Azuma (Ratiu and Azuma, 2015), presenting an inconsistent effect of bilingualism on cognitive performance. This second view holds that because working memory in bilinguals is occupied by more information, their performance would be hindered.

Due to the inconsistency mentioned above, this study aims to answer these two questions: 1. Are bilinguals more resilient than monolinguals when faced with high cognitive load? 2. Does the amount of cognitive load matter to differentiate between bilinguals and monolinguals’ cognitive performance? To this end, we recruited a group of Persian-English late bilinguals (N=19) by purposive sampling and, an age and education matched group of Persian monolinguals (N=21) as the control group. We compared their behavioral performances (response accuracy and reaction times) in a dual interference task (color-word Stroop task with different levels of difficulty which impose different levels of cognitive load).

### The rationale for the selection of the task

Developed by John Ridley Stroop, Stroop task is concerned with three theories, including selective attention, speed of processing theory, and cognitive load theory (Lavie and Viding, 2004). Therefore, it has extensively been used to measure cognitive performance. Using color-word Stroop tasks will help us to measure the intrinsic cognitive load. Adding a concurrent verbal task to the Stroop task will allow us to measure the extraneous cognitive load. A quite similar approach where the authors asked participants to perform a verbal production, information search or learning tasks concurrently with a cognitive task that has been used in a number of studies (Gwizdka, 2010; Hazan-Liran and Miller, 2017; van der Heiden, R. M., Janssen et al., 2020). This method is promising to measure intrinsic and extraneous cognitive load.

Besides performing the computerized Color-Word Stroop task while totally attending to the task, we have selected two verbal tasks with different difficulty levels to be performed simultaneously with the Stroop task; alphabet reciting, which is considered as an automatic verbal production process, and counting odd numbers, which is cognitively more demanding. This leads to three levels of task difficulty and thus three levels of cognitive load: 1)The color-word Stroop task performed in silence, 2) the color-word Stroop task while alphabet reciting, and 3 the color-word Stroop task while counting odd numbers. This would investigate the intrinsic and extraneous loads imposed on the participants. Extraneous load emanates from processing task-irrelevant information (Hazan-Liran and Miller, 2017), in our design, alphabet reciting and counting odd numbers while doing the Stroop task, and intrinsic load is caused by the inherent demands and difficulties of the Stroop task itself (Paas et al., 2003). Importantly, the stability of participants’ performance in the Stroop task with an increase in cognitive load can be considered as an objective measure of cognitive resilience (see for example: Shields et al., 2017)). To our best knowledge, this is the first study to compare cognitive performances in bilinguals and monolinguals considering different levels of task difficulty.

## Method

### Participants

Bilingual group: Nineteen bilingual speakers, N= 5 males, with Persian as their mother tongue (L1) and English as their second language (L2) aged between 20 to 34 (26.3 ±3.6), were recruited.

Monolingual group: A group of 21 Persian monolingual participants, aged between 24-35 (27.8 ±3.1), N= 9 males, were included in the monolingual group of the study.

All participants of both groups were right-handed. They had not reported any history of neurological, psychiatric or other medical conditions that could affect our study. They all had normal or corrected to normal vision. The two groups were matched for age and education. Voluntary participation was ensured using a signed consent form in accordance with the Declaration of Helsinki. The protocol of the study was approved by the local ethical committee at the School of Cognitive Sciences, IPM, Iran.

### Questionnaire on psychological resilience

A Persian standardized version of the Conner-Davidson Resilience Scale (CD-RISK) was administered to provide a subjective measure of individuals’ psychological resilience in both groups. This would allow us to control for effects of psychological resilience on participants’ cognitive resilience. CD-RISK measures the ability to cope with stressful situations and adversity. Respondents rate 25 different items on a scale from 0 (“not true at all”) to 4 (“true nearly all the time”) (Khoshouei, 2009).

### Second language evaluation

#### Receptive language evaluation

The vocabulary subtest was taken from the computer-based DIALANG language diagnosis system and administered to evaluate vocabulary knowledge in English (Zhang and Thompson, 2004). In this test, participants indicate for each of 75 stimuli whether it is a correct word in English or a highly word-like pseudo-word. The DIALANG score between 600 and 900 indicates advanced proficiency and a score between 400 and 600 indicates good basic vocabulary. Bilingual participants that were included for the analyses in the present study had an average DIALANG score of 596.2, while monolingual participants had a mean DIALANG score of 281.5.

#### Productive language evaluation

Productive Vocabulary Levels Test (PVLT) (Laufer and Nation, 1999) was used to evaluate English productive vocabulary in both monolingual and bilingual participants. The PVLT samples 18 items at five different word-frequency levels. According to Nation and Waring (1997), second language learners with knowledge of the most frequent words will know around 80% of the running words in a written or spoken text (Nation and Waring, 1997). Low-frequency words cover the remaining 20%. According to their performance in the PVLT, bilingual participants that were included for the analyses in the present study had knowledge of the most frequent words of at least 73% and knowledge of low-frequency words of at least 48%. Monolingual participants had very limited knowledge of both high and low-frequency words (31% and 10%, respectively).

#### Language use questionnaire

Participants from both monolingual and bilingual groups filled out the language experiences and proficiency questionnaire (LEAPQ) (Marian et al., 2007). Age of acquisition, immersion, and self-evaluation of language use were assessed using questions regarding different contexts of language use and exposure (i.e., with friends, family, watching TV/listening to the radio, reading books). The participants listed all the languages they use or have exposure to in daily basis.

The use of this questionnaire was essential to ensure the monolingualism of our control group. In the language use questionnaire, monolinguals indicated that their language use and exposure to Persian is 90%. While in the bilingual group, around 40% of their daily language usage and exposure (reading a book, watching movies, communication with friends) were in English. Table 1 shows the data of L2 usage and proficiency of the participants.

**Table 1.** Second language proficiency scores and usage

| **Variable** | **Bilinguals** |  | **Monolinguals** |  |
| --- | --- | --- | --- | --- |
|  | **Mean** | **SD** | **Mean** | **SD** |
| **L2 Test** |  |  |  |  |
| DIALANG score | 596.15 | 101 | 281.5 | 116 |
| PVLT high frequency words (%) | 73 | 0.1 | 48 | 0.1 |
| PVLT low frequency words (%) | 31 | 0.1 | 10 | 0.05 |
| PVLT total scores (%) | 54 | 0.1 | 24 | 0.1 |
| **Use of L1** |  |  |  |  |
| Work/study (%) | 57 | 12 | 89 | 9 |
| TV/Radio | 61 | 11.2 | 92 | 5 |
| **Use of L2** |  |  |  |  |
| Work/study (%) | 35 | 7 | 9 | 6 |
| TV/Radio | 41 | 8 | 6 | 5 |

### Study procedure

All participants performed three blocks of a color-word Stroop task with three different levels of task difficulty; Difficulty level 1: Simple Stroop task, difficulty level 2: Stroop Task and alphabet reciting, and difficulty level 3: Stroop Task and odd numbers counting. The order of tasks was counterbalanced across participants (Figure 1-A).

**Figure 1.**
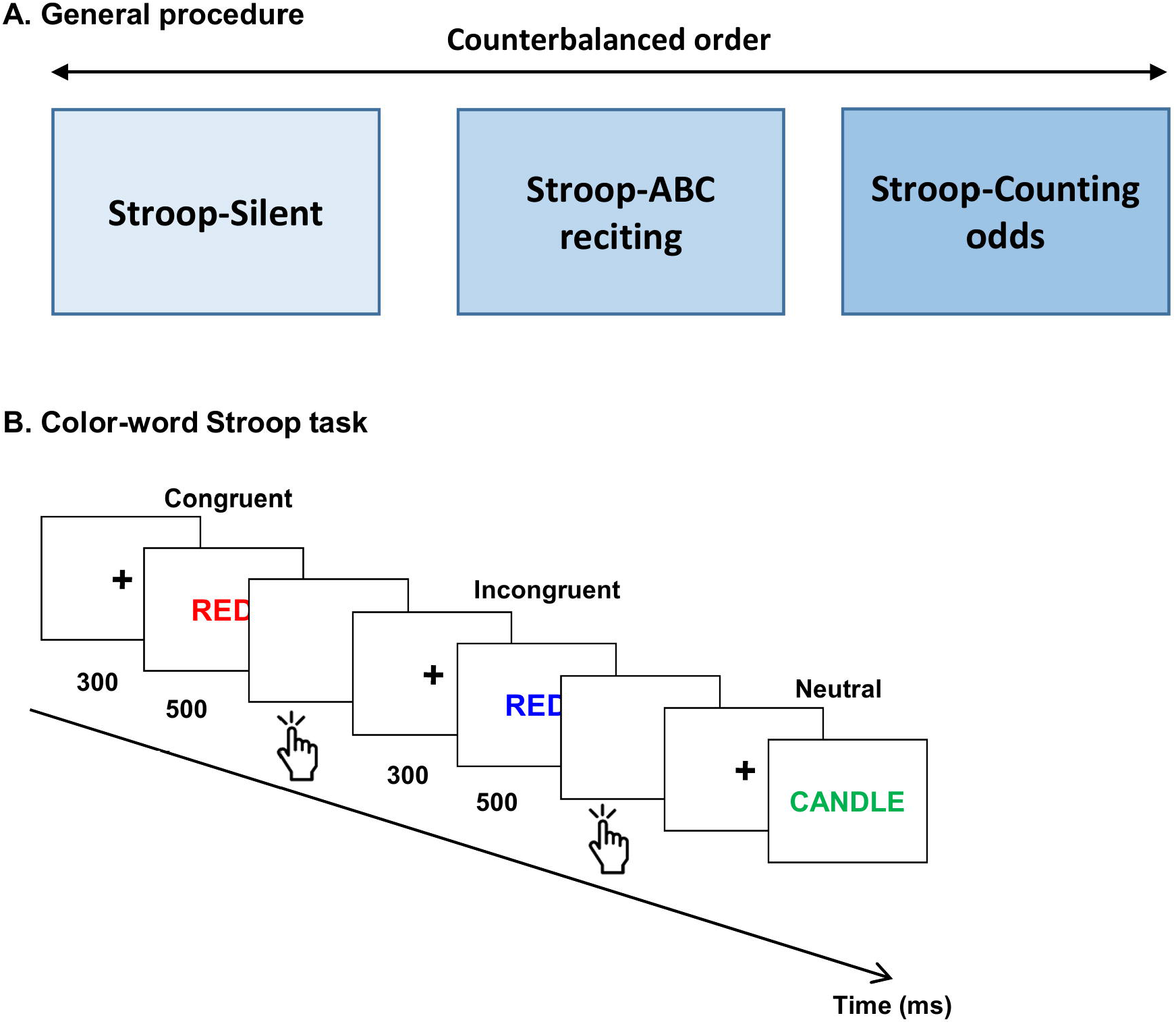
Study procedure and tasks. (A) Study procedure: each participant was asked to do the Stroop task with three levels of difficulty: 1. Stroop task in Silence, 2. Stroop task while alphabet reciting and 3. Stroop task while counting odd numbers. The order of these tasks was counterbalanced across participants. (B) Color-Word Stroop task: Three types of stimuli were used including Congruent, Incongruent, and Neutral. Participants were instructed to select the printed color of each word (i.e., red, green, blue, or yellow) on the keyboard; Participants were asked to press ‘b’ for blue, ‘r’ for red, ‘y’ for yellow, and ‘g’ for green as fast as possible.

#### Dual-task interference

##### Difficulty level 1: Simple Stroop Task

Participants performed a modified version of the color-word Stroop task used by (Andrews-Hanna et al., 2011). Three types of stimuli were used including Congruent, Incongruent, and Neutral. As in a traditional Stroop task, participants select the color in which a word is printed across two main trial types: Congruent (i.e., red shown in red text) and Incongruent (i.e., red shown in green text). In addition, Neutral trials are presented in which the stimulus is a color-neutral word (e.g., chair) shown in colored text. Participants were instructed to select the printed color of each word (i.e., red, green, blue, or yellow) on the keyboard; Participants were asked to press ‘b’ for blue, ‘r’ for red, ‘y’ for yellow, and ‘g’ for green as fast as possible. Each trial starts with a fixation cross lasting for 300ms. Then, a target word appears on the screen for 500ms. The participants were asked to respond in order to go to the next trial. Six blocks of 90 trials (with the same proportion of the three types of stimuli) with a random presentation of Congruent, Incongruent, and Neutral trials were recorded. There was a 2 minutes rest between the two blocks. The inclusion of the neutral stimuli across all blocks reduces potential habituation effects and prevents participants from merely reading the words during the Congruent trials. The neutral trials included a list of four high-frequency (basket, scarf, candle, & photo) (Figure 1-B).

##### Difficulty level 2: Stroop Task and alphabet reciting

In this step, subjects performed the same Stroop task (the same details in the previous part) while reciting the alphabet simultaneously.

##### Difficulty level 3: Stroop Task and odd numbers counting

At this stage, subjects performed the same Stroop task (the same details in the previous parts) while counting odd numbers simultaneously.

At the beginning of the experiment, a short training run was performed to assure that the subjects were ready to do the main task.

### Behavioral data analyses

The “Stroop effects” were calculated as reaction time to incongruent - reaction time to congruent trials of the three conditions (task difficulty levels 1, 2, and 3).

Similarly, the difference of accuracy performance (i.e., “Δ accuracy”: accuracy in congruent - accuracy in incongruent trials) was measured for task difficulty levels 1, 2, and 3.

The Stroop effect and Δ accuracy scores were implemented in a mixed between-within design ANOVA with between factor Group (Bilingual, Monolingual), and within factor Difficulty Level (1, 2, 3).

## Results

### Observations on the verbal production

Verbal production (alphabet reciting, and counting odd numbers) was qualitatively evaluated by the examiner. Slow-down and short stop periods were systematically observed at the moment the participant was presented by the stimulus and a keypress to respond was required.

### Psychological resilience score

No significant difference between the two groups was found in the resilience score using CD-RISK; independent sample t-test p=0.13.

### Dual-task interference behavioral results

Mixed design ANOVA analyses on Stroop effect, revealed main effects of Difficulty Level (F(2,76) = 12.7, p < 0.001, η2p = 0.25), main effect of Group (F(1,38) = 7.18, p = 0.01, η2p = 0.159). No significant interaction was found between Group and Difficulty Level (F(2,76) = 1.52, p = 0.22, η2p = 0.04) (Figure 2-A).

**Figure 2.**
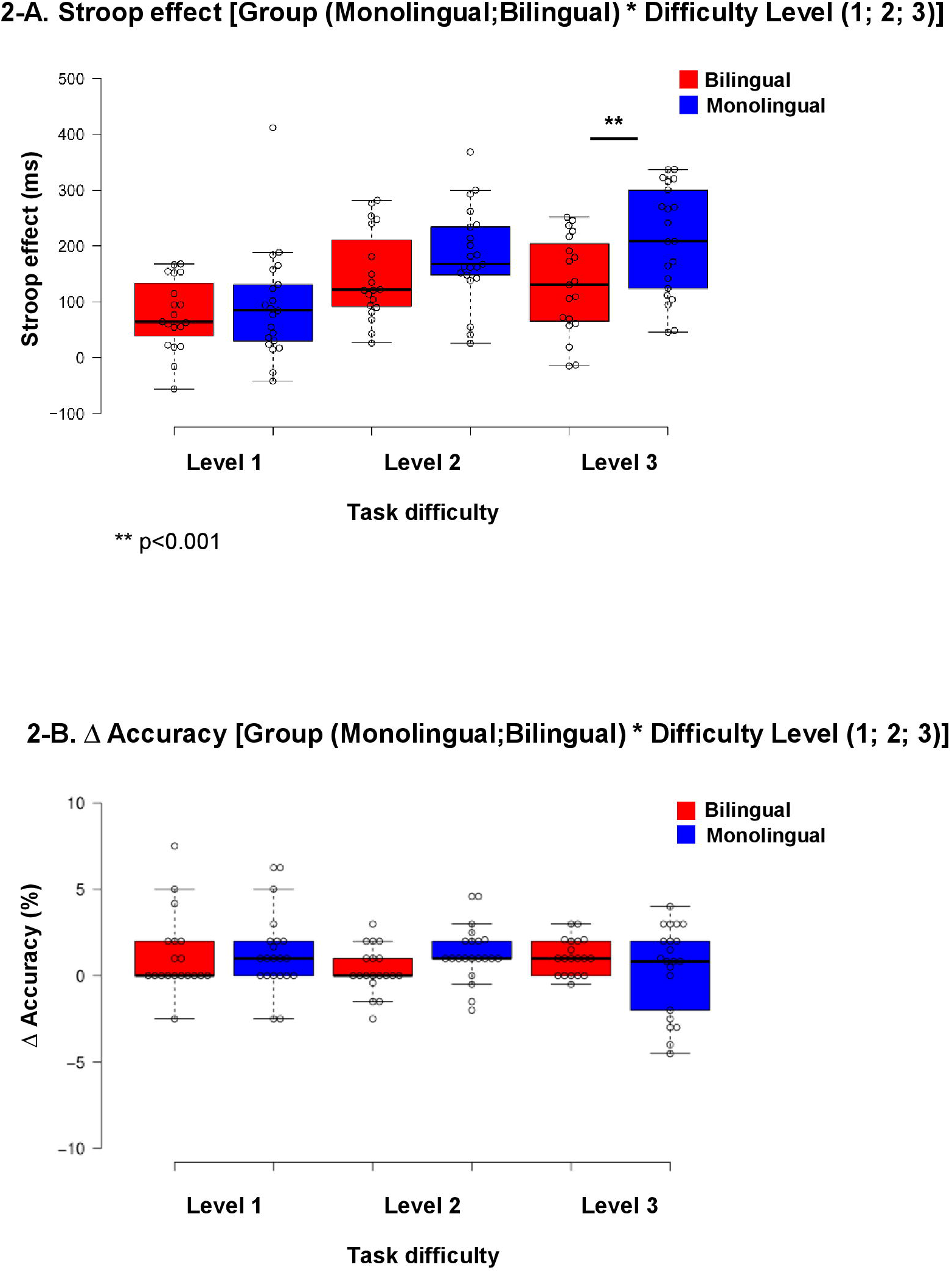
Behavioral results. (A) Stroop effect: Main effects of Difficulty and main effect of Group were found. No significant interaction was found between Group and Difficulty Level. (B) Δ Accuracy: No main effects of Difficulty Level or Group, nor interaction between these factors was seen.

Mixed design ANOVA analyses on Δ accuracy, showed no main effects of Difficulty Level and Group (F (2,76) = 0.92, p = 0.40, η2p = 0.02 and F(1,38) = 0.14, p = 0.70, η2p = 0.004 respectively), nor interaction between these factors (F(2,76) = 0.62, p = 0.54, η2p = 0.02) (Figure 2-B).

## Discussion

This study aimed to investigate whether bilinguals are more resilient than monolinguals when faced with high cognitive load, and, whether the amount of cognitive load affects these two groups’ cognitive performance differently. We measured participants’ performance using the Stroop effect (the reaction time differences in congruent and incongruent trials), and Δ accuracy (the accuracy differences in congruent and incongruent trials) in response to low, medium, and high task difficulty levels. These difficulty levels were defined and were related to the level of the extraneous cognitive load imposed on the participants using different verbal production tasks which were done concurrently with a color-word Stroop task (i.e., Level1: Silence, Level2: alphabet reciting, Level 3: counting odd numbers). In levels 1 and 2, considered as simple and moderate, the Stroop Effect of both groups was proved to be the comparable. The results have shown that the Stroop effect does not increase significantly in bilingual participants when task difficulty level increases. While in the monolingual group, an increase in cognitive load resulted in a larger Stroop effect. Therefore, the interference effect of the difficult conditions was reported to be lower for the bilinguals rather than monolinguals.

This finding suggests that bilingual participants show stability in their cognitive performance, and have therefore a higher resilience when faced with high cognitive load. According to Sweller (Sweller, 1988), cognitive overload occurs when the demand for processing the task exceeds the attentional resources and cognitive capacity. The research confirms the facilitating effect of bilingualism on attention resources and cognitive capacity in this regard. The explanation can be derived from the data that bilinguals are better able to avoid the interferences in high demand cognitive tasks than monolinguals. In other words, by managing two different processes, bilingualism will result in a more efficient use of the attention resources and cognitive ability.

Our results are in line with a great body of previous research suggesting that bilingual speakers outperform monolinguals in domain-general cognitive control. Bialystok (Bialystok et al., 2005, 2012) notes that bilingualism is one of the factors affecting cognitive capacity and cognitive function to some degree. This effect can be altered by the age of acquisition and the proficiency level (Blom et al., 2017; Hernandez et al., 2012; Kousaie and Phillips, 2011; Marian, V., & Shook, 2012; Mechelli et al., 2004). Coderre, van Heuven, and Conklin also evaluated the Stroop effect in bilinguals and their results was in favor of the bilingualism advantage. They explained this outcome through the critical position of language proficiency and immersion (Coderre et al., 2013). Later on, Coderre and van Heuven used electrophysiological approach and applying Stroop task, and found that bilinguals have better performance in executive control tasks than the monolinguals. The authors explained that bilinguals have a better system for handling interferences (Coderre and van Heuven, 2014). In addition, Yang and Yang have shown that bilingualism facilitates control processing which is the primary function of working memory and this is justified by the demand imposed on attentional control in working memory (Yang and Yang, 2017). Working memory is influenced by bilingualism through retrieving the intended element from the lexicon of the two systems and the inhibition ability in the task-irrelevant stimuli (Nguyen and Astington, 2014; Soveri et al., 2011).

The approach used in our study to investigate the cognitive resilience in conditions with different cognitive load was based on the recommendation from previous works; as such, Simon and colleagues attempted to investigate executive function in terms of increasing cognitive load. The authors found that the output of the participants would be impeded by increasing the working memory load (Simon et al., 2016). Similarly, Shields et al., investigated cognitive resilience in the student-athletes and a control group using the Stroop effect. They found that cognitive resilience among student-athletes resulted in psychological adaptation over the course of a season (Shields et al., 2017). Besides, the key consideration in the analyses of the Stroop test is the demonstration of resistance to task interference and information incongruency (Block, 2004).

In addition to previous methods, the current study improved the design by adding different difficulty levels to the Stroop task. This is specifically noteworthy when faced with controversial findings regarding bilingual advantage on executive functions. Importantly, several studies put into question the positive effect of bilingualism on executive performance (e.g., Paap and Greenberg, 2013; Ratiu and Azuma, 2015). Most of the previous negative results regarding bilingual benefits on cognitive performance come from the studies in which a single level of task difficulty has been used to compare bilingual subjects’ executive performance with monolinguals’ (See for example: Dunabeitia et al., 2013; Paap and Greenberg, 2013). Furthermore, Yang and Yang demonstrated that, while bilinguals outperformed monolinguals in the attention impeded Stroop-span task, monolinguals and bilinguals had the same performance in the operation span and Stroop-span tasks (Yang and Yang, 2017). To illustrate more tangibly, the performance of bilinguals and monolinguals was similar in the tasks absent from interference. To conclude, the findings of the current study and related literature indicate that no distinction can be made between bilinguals and monolinguals cognitive resilience in the condition with a lower degree of cognitive load.

Considering figure 2, an absence of difference between monolinguals and bilinguals performance in Stroop task (as reflected in the Stroop effect) in the simple and moderately difficult task could somehow explain the inconsistent pattern of results of the previous studies; according to Paas and colleagues, the basic low-level interactive conditions can be processed with a low-level application of cognitive resources and pre-stored schemas (Paas et al., 2010). On the other hand, performing more complex tasks requires a higher level of interactivity. Therefore, in difficult conditions, the difference between those who have a higher cognitive ability and lower cognitive ability is highlighted. Ultimately, the contrasting performance of bilinguals and monolinguals showed the supporting impact of bilingualism in high cognitive load tasks.

It is noteworthy that the obtained results of the present study are not explained by any possible difference in psychological resilience between the two groups. This was ensured using a CD-RISK questionnaire on resilience which resulted in comparable scores for both groups. Although controversial, a possible interaction between psychological resilience and cognitive performance has been investigated in a number of previous studies (e.g., Parsons et al., 2016; Shields et al., 2017). One explanation for the unexpected outcome can be explained by the invalidity of certain contexts of the self-report inventory based on social desirability bias (Demetriou et al., 2015).

In contrast to our results on the Stroop effect based on reaction time, regarding Δ accuracy - as defined by the difference between response accuracy in congruent and incongruent trials in our study-, no significant difference between monolingual and bilingual groups was observed with an increase in cognitive load. This could be explained in accordance with the results of previous works that suggest that response accuracy is not sensitive enough to measure the participants’ performances when considered independent from response time (Kousaie and Phillips, 2011).

Due to the fact that the pressure on people has increased more significantly than before, the cognitive load has been investigated in different domains. According to cognitive load theory, every task requires a continuum of interactivity. The easier one needs simple processing in terms of automaticity, while the complex task requires more complex processes to be performed (Sweller et al., 2019). Cognitive load theory takes the notion of a schema that moves from long term memory to the working memory and vice versa, and it features the cognitive capacity in the state of being inundated with adverse information.

The current study sheds more light on the nature and importance of bilingual advantage. This study experimentally investigated the cognitive resilience ability in the tasks with different cognitive load. However, the results of this study are limited to behavioral level and further research including neuroimaging techniques would add more value to resolve the controversies in the literature.

## Conclusion

In summary, behavioral measures of our study showed that bilinguals could better monitor the discrepancies and keep a steadier cognitive performance than monolinguals when task-irrelevant demand exists. Although, it is notable that the degree of the cognitive load should also be taken into consideration. Because, as demonstrated in our results, easy to moderate conditions would not differentiate cognitive control differences of bilinguals and monolinguals. These findings support the view of bilingual advantage on cognitive performance. However, our results emphasize that such cognitive advantage is mostly seen when cognitive demand increases.

## Acknowledgements

This study was supported by internal funding from the School of Cognitive Sciences, Institute for Research in Fundamental Sciences, Iran.

## Authors’ contribution

S.D. contributed to the conception, data collection, and writing of the manuscript.

Z. S. contributed to the conception, data analyses, and writing of the manuscript.

N.R. contributed to the conception, data analyses, and writing of the manuscript.

